# Genomic detection of a secondary family burial in a single jar coffin in early Medieval Korea

**DOI:** 10.1101/2022.05.09.491093

**Authors:** Don-Nyeong Lee, Chae Lin Jeon, Jiwon Kang, Marta Burri, Johannes Krause, Eun Jin Woo, Choongwon Jeong

**Affiliations:** School of Biological Sciences, Seoul National University, Seoul, 08826, Republic of Korea; Department of Anthropology, Seoul National University, Seoul, 08826, Republic of Korea; Central Institute of Cultural Heritage, Daejeon, 34029, Republic of Korea; Max Planck Institute for the Science of Human History, Jena, 07745, Germany; Max Planck Institute for Evolutionary Anthropology, Leipzig, 04103, Germany; Department of History, Sejong University, Seoul, 05006, Republic of Korea

**Keywords:** ancient DNA, family burial, jar coffin, Gunsan Dangbuk-ri, population genomics

## Abstract

**Objectives:** Family relationship is a key to understand the structure of past societies but its archaeological reconstruction mostly stays circumstantial. Archaeogenetic information, especially genome-wide data, provide an objective approach to accurately reconstruct the familial relationship of ancient individuals, thus allowing a robust test of an archaeology-driven hypothesis of kinship. In this study, we applied this approach to disentangle the genetic relationship of early Medieval individuals from Korea, who were secondarily co-buried in a single jar coffin.

**Materials and Methods:** We obtained genome-wide data of six early Medieval Korean individuals from a jar coffin. We inferred the genetic relatedness between these individuals and characterized their genetic profiles using well-established population genetics methods.

**Results:** Congruent with the unusual pattern of multiple individuals in a single jar coffin, genome-wide analysis of these individuals shows that they form an extended family, including a couple, their two children and both paternal and maternal relatives. We show that these early Medieval Koreans have a genetic profile similar to present-day Koreans.

**Discussion:** We show that an unusual case of a secondary multiple burial in a single jar coffin reflects family relationship among the co-buried individuals. We find both paternal and maternal relatives co-buried with the nuclear family, which may suggest a family structure with limited gender bias. We find the genetic profile of early Medieval Koreans similar to that of present-day Koreans, suggesting no substantial genetic shift in the Korean peninsula for the last 1,500 years.

**Research Highlights:**

- Ancient genome-wide data find a family buried together in a jar coffin in early Medieval Korea.
- These early Medieval Koreans have a genetic profile similar to present-day Koreans.

## Introduction

The burial practice using jar coffins was widespread throughout past societies in East Asia including Southeast Asia, northern China, Manchuria, the Japanese archipelago, and the Korean Peninsula, starting from the Neolithic period and continuing to the historic times (Bacvarov, 2006; Boeyens et al., 2009; W. Kim, 1973; Shewan et al., 2020). In the Korean peninsula, while prehistoric and historic burials with jar coffins are frequently found in all regions, they are most concentrated in the southwestern Korea, such as Naju and Yeongam regions, during the 4-6^th^ centuries AD (E.-J. Kim, 2021; E. K. Kim, 2020; M. J. Kim et al., 2010; Oh, 2008; Park, 2010). Although burials with large, human-height-scale jar coffins are famous, most jar coffins found in Korea are much smaller than human body. Mostly, these small jar coffins host a single individual’s skeleton secondarily collected after cremation. The practice of cremation and the following secondary burial often result in a loss of some skeletal elements, destruction of detailed morphological features that convey information on the sex, age, and pathological history of the individual, and poor preservation of biological macromolecules such as proteins and nucleic acids. For this reason, skeletal remains from the jar coffins have not been actively investigated in bioanthropological and archaeogenetic studies, leaving the characteristics of people buried in the jar-coffins an open question.

The Dangbuk-ri archaeological site is located at Gunsan city in the west coastal region of South Korea (Figure 1). Excavated during Summer 2016, this site includes several burials of the early Medieval period (5-7^th^ century AD; or the “three kingdoms” period). Among the various types of burials is one jar coffin burial housing skeletal remains of multiple individuals (Figure 2). Considering the burial context, it is most likely that the skeletal elements of these individuals were collected secondarily into this jar coffin and buried at the same time. This case is considered unusual because most jar coffin burials house only one individual and because individuals in the other types of multiple burials in the early Medieval Korea were usually added into the burial in a sequential manner over an extended period of time (E.-J. Kim, 2021).

**Figure 1.**
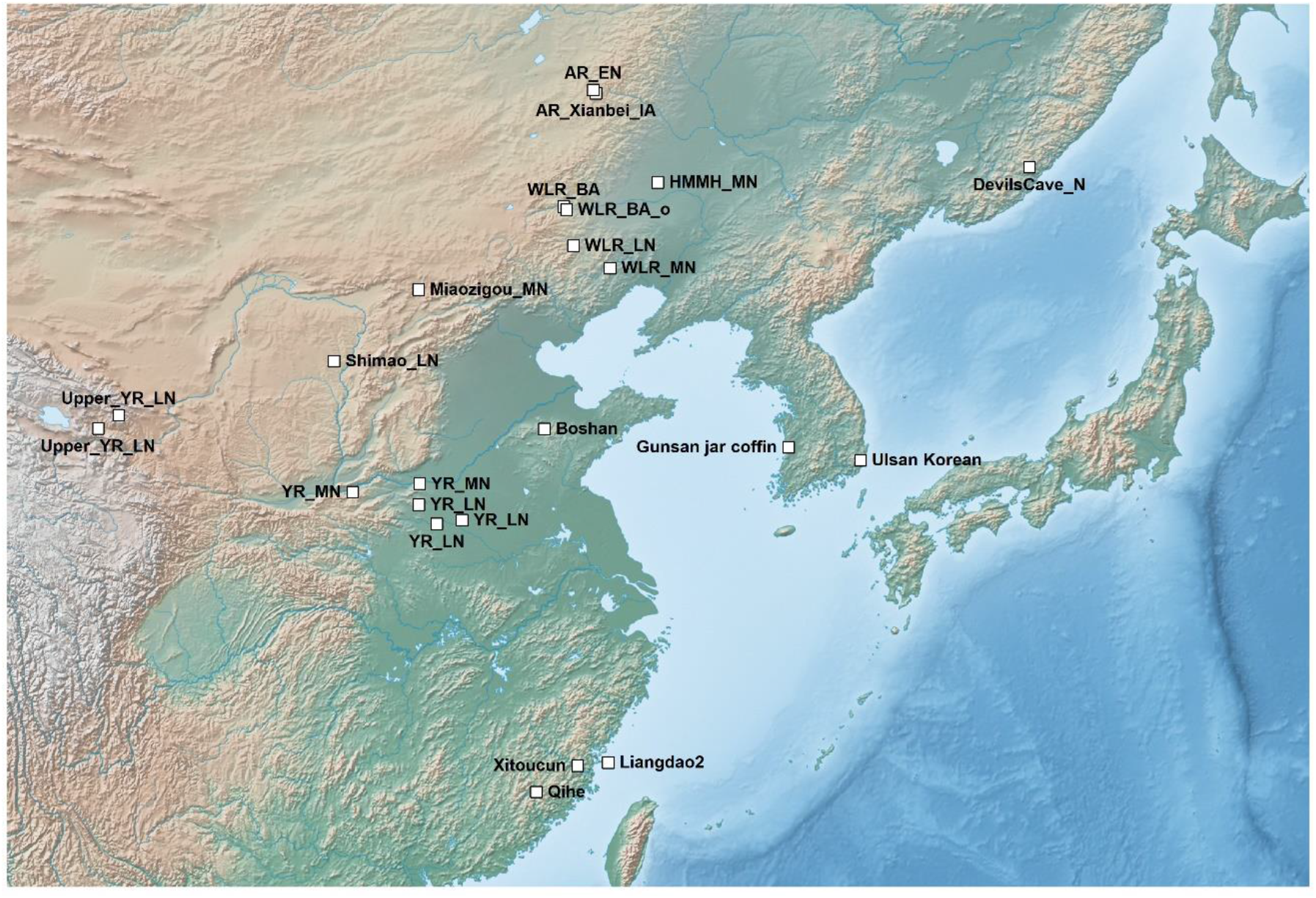
The geographic location of the key ancient and present-day populations used in this study. Except for present-day Ulsan Koreans from southeastern Korea, all the other groups marked on the map represent ancient East Asian groups. The base map is produced using the Natural Earth public domain map dataset (https://www.naturalearthdata.com/downloads/10m-raster-data/10m-cross-blend-hypso/). AR_EN = Amur River Early Neolithic individuals from the Zhalainuoer/Wuqi site; AR_Xianbei_IA = Amur River Iron Age individuals from the Xianbei context of the Mogushan site; Boshan = an early Neolithic individual from Shandong region; DevilsCave_N = Early Neolithic individuals from the Russian Far East; HMMH_MN = Middle Neolithic individual from the Haminmangha site; Liangdao2 = an Early Neolithic individual from the Liangdao site; Miaozigou_MN = Middle Neolithic individuals from the Miaogizou site; Qihe = Early Neolithic individuals from the Qihe site; Shimao_LN = Late Neolithic individuals from the Shengedaliang site; Upper_YR_LN = Upper Yellow River Late Neolithic individuals from the Jinchankou and Lajia sites; WLR_MN = West Liao River Middle Neolithic individuals from the Banlashan site; WLR_LN = WLR Late Neolithic individuals from the Erdaojingzi site; WLR_BA/WLR_BA_o = WLR Bronze Age individuals from the Longtoushan site; Xitoucun = Late Neolithic individuals from the Xitoucun site; YR_MN = Yellow River Middle Neolithic individuals from the Xiaowu and Wanggou site; YR_LN = YR Late Neolithic individuals from the Haojiatai/Pingliangtai/Wadian site.

**Figure 2.**
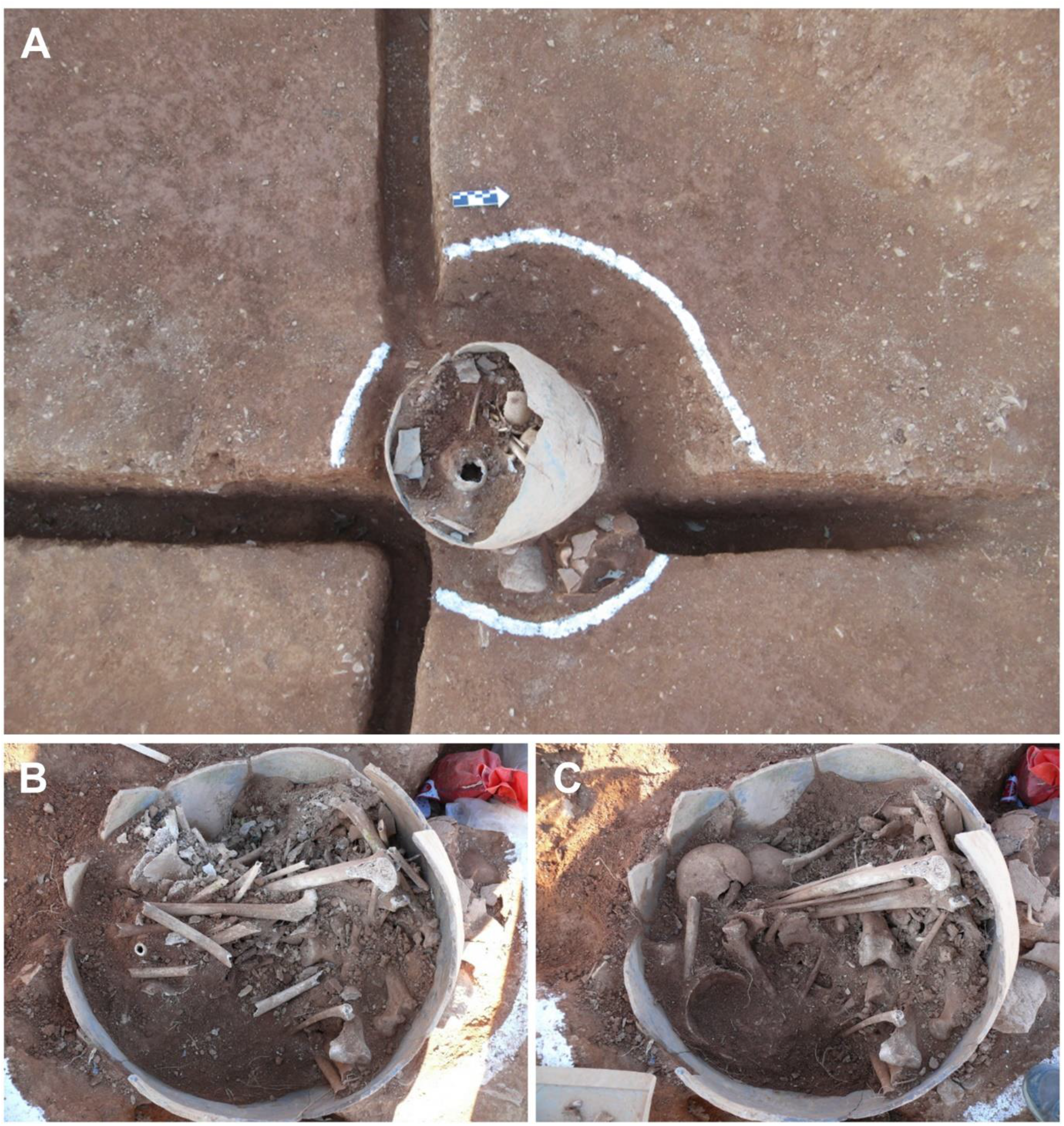
Revealed skeletons in a jar-coffin, Dangbuk-ri site *in situ.* (A) A wide view of the Gunsan jar-coffin. (B, C) The remains of the Gunsan jar-coffin individuals.

In this study, we investigated the genetic relatedness between these co-buried individuals and their genetic profiles using genome-wide data. In addition to revealing the familial relationship among them, we model their genetic profiles in terms of their relationship with ancient and present-day East Asian populations, including present-day Koreans, in high-resolution by utilizing rich information in genome-wide data.

## Materials and Methods

### Skeletal analysis

All skeletons were macroscopically examined by the two authors (E.J.W and C.L.J). In order to estimate the minimum number of individuals (MNI) in a jar, fragments were first identified by element type, and conjoining process was conducted to refit fragments from the same bone. Identified elements were classified as either adult or subadult according to the degree of dental development, long-bone epiphyseal fusion, diaphyseal length, cranial size and cortical thickness. Bones with fully fused epiphyses and within the adult-sized range elements were classified as adults. Meanwhile, skeletal elements with unfused epiphyses and within the size range of the subadults were classified into the subadult category. After the skeletal elements were sorted to individuals, age estimation was further refined based on morphological features. The value of MNI was derived by sorting elements into lefts and rights, and then taking the greatest number as the final estimate following a published protocol (T. E. White, 1953). Severely fragmented skeletal elements such as ribs, vertebrae and phalanges were excluded from calculating MNI due to uncertainty in siding. Then, visual pair-matching was conducted to decide if principal limb bones were from a single individual through the comparison of right and left elements. Finally, different parts of the skeletal elements were segregated into individuals based on the examination of degenerative changes, articulation, robusticity, age, and sex, along with osteometric sorting (Adams & Byrd, 2014).

The sex and age were estimated for each skeletal individual, based on the Standards for Data Collection (Buikstra & Ubelaker, 1994; Ubelaker, 1999). Sex was estimated using morphological features of the skull and robusticity of limb bones. We were unable to use the pelvis for sex estimation because most pelvic elements in the jar-coffin were severely fragmented. The age at death of each individual was estimated by dental attrition, number of antemortem tooth loss, cranial suture closure, degeneration of the auricular surface and pubic symphyseal surface, and degenerative changes of joint in limb bones and the vertebral column. The age of subadults was estimated using the degrees of tooth formation and eruption, epiphyseal fusion or diaphyseal length (Buikstra & Ubelaker, 1994; T. D. White & Folkens, 2005).

### Sampling of skeletal materials

Sample selection was performed by E.J.W. and C.L.J. in the Bioanthropology laboratory (Department of Anthropology, Seoul National University). As compact bones are preferred for ancient DNA analysis, intact petrous parts of the temporal bone from seven individuals were chosen. Two out of nine individuals were not included in the analysis since the temporal parts including petrous were not well preserved. In case of five adults, the left petrous bone was selected, for two subadults, the right petrous parts were sampled. The outer surface of the petrous pyramid inside the skull was ground with a 4.8 mm cutter burr attached to a Dremel 9100-21 Fortiflex 2.5-Amp Stationary Flex Shaft Precision Rotary Tool (Dremel, Mount Prospect, IL). Then the inferior border of the cochlea was ground to create a small opening into the osseous labyrinth (Sirak et al., 2017). Into this opening, a 3.2 mm engraving cutting burr was applied in a circular motion to obtain bone powder. The powder was collected in a sterilized paper foil and placed in a sterile 1.5 ml Eppendorf tube for DNA extraction.

### Ancient DNA laboratory work and sequencing

For each of the seven individuals, a double-strand double-indexed Illumina sequencing library was built from metagenomics DNA extracted from 30-50 mg of bone powder. DNA extraction and library preparation were performed using previously published protocols (Dabney et al., 2013). For library preparation, a partial treatment of the uracil-DNA-glycosylase (UDG) enzyme was included following a published protocol (Rohland et al., 2015) to confine deaminated bases to the ends of the reads. All laboratory works up to library prepraration step were performed in a dedicated ancient DNA (aDNA) clean room facility of the Max Planck Institute for the Science of Human History (MPI-SHH), Jena, Germany. For six of the seven individuals with sufficient levels of human DNA preservation, ranging 0.1-2.6%, two rounds of in-solution capture for 1.24 million ancestry-informative single nucleotide polymorphisms (SNPs) (the “1240K” panel) was performed to enrich libraries (Mathieson et al., 2015). All libraries were sequenced using single-end 76 base pair (bp) sequencing on the Illumina HiSeq 4000 platform following the manufacturer’s protocols.

### Ancient DNA sequencing data processing and authentication

aDNA sequencing data were processed using the EAGER v1.92.50 wrapper (Peltzer et al., 2016). Within the EAGER wrapper, Illumina adapter sequences were first trimmed from raw reads using AdapterRemoval v2.3.0 (Schubert et al., 2016). Adapter-trimmed reads of 30 bp or longer were mapped to the human reference genome with decoy sequences (hs37d5) using the aln/samse modules in BWA v0.7.17 with “-n 0.01” option (Li & Durbin, 2009). PCR duplicates were removed using DeDup v0.12.5 (Peltzer et al., 2016), assuming that both ends of the reads were known. Unique mapped reads with Phred-scaled mapping quality score 30 or higher were kept using samtools v1.9 (Li et al., 2009). To remove deamination-based misincorporations (5 ‘ C>T and 3 ‘ G>A), the first and last two bases of each read were soft-masked using the trimBam module of bamUtils v1.0.14 (Jun et al., 2015). Finally, pseudo-haploid genotype data were determined by randomly sampling a single high-quality base (Phred-scaled base quality score 30 or higher) per site per individual using samtools mpileup and pileupCaller v1.4.0.5 (downloaded from https://github.com/stschiff/sequenceTools). For C/T and G/A SNPs, end-masked BAM files were used. For the remaining SNPs that are not affected by post-mortem deamination, BAM files without end-masking were used to maximize sequence data usage. The authenticity of sequence data was checked by multiple measures. First, chemical modifications typical of aDNA molecules were tabulated using mapDamage v2.0.9 (Jonsson et al., 2013). Second, mitochondrial DNA contamination was estimated using schmutzi v1.5.5.5 (Renaud et al., 2015). Third, X chromosome-based estimation of nuclear DNA contamination was performed for four male individuals using the contamination module of the ANGSD v0.929 program (Korneliussen et al., 2014). Mitochondrial haplogroups were determined by applying the HaploGrep v2 program (Weissensteiner et al., 2016) to the consensus sequences called by the log2fasta program in schmutzi with “-q10” filter. Y haplogroups were determined using a modified version of the yHaplo program with “--ancStopThresh 10” option to prevent the root-to-tip search from halting in an internal branch due to missing data (downloaded from https://github.com/alexhbnr/yhaplo) (Poznik, 2016).

### Reprocessing of whole genome sequences of present-day Koreans

We downloaded high-coverage whole genome sequencing data of 104 Koreans from the Ulsan city from KoVariome data (ftp://biodisk.org/Release/KPGP/KPGP_Data_2017_Release_Candidate/). For the FASTQ files with the Phred-64 scale quality scores, we rescaled quality scores to the Phred-33 scale using seqtk v1.3-r106 with “seqtk seq –VQ64” command (https://github.com/lh3/seqtk). We aligned reads to the human reference genome (hs37d5) using the BWA mem program v0.7.17 with “-M” flag (Li & Durbin, 2010). We kept properly aligned paired-end reads by applying “-f 0×0003” filter in samtools view v1.9 (Li et al., 2009) and merged per-lane BAM files into per-individual ones using samtools merge. We removed duplicates using Picard MarkDuplicates v2.20.0 (downloaded from https://broadinstitute.github.io/picard/) and kept reads with Phred-scaled mapping quality score 25 or higher. From these analysis-ready BAM files, we produced two genotype calls for the 1,233,013 SNPs in the 1240K panel. First, we calculated genotype likelihoods for each individual using the UnifiedGenotyper module of the Genome Analysis ToolKit (GATK) v3.8.1.0 (McKenna et al., 2010) with the “--allSitesPLs” flag, calculated posterior genotype probability by multiplying genotype likelihoods with the GATK default prior [0.9985, 0.0010, 0.0005], and took genotype calls with posterior probability 0.900 or higher. Second, we randomly sampled a base with the Phred-scaled base quality score 30 or higher from a read with mapping quality score 30 or higher per site per individual to create a pseudo-haploid call that mimics the genotype calling procedure of low-coverage ancient individuals. For this procedure, we used samtools mpileup and pileupCaller v1.4.0.5. We merged the two genotype calls of present-day Ulsan Koreans with the genome-wide data of world-wide present-day populations typed on the Affymetrix Axiom^®^ Genome-Wide Human Origins 1 array (“HumanOrigins”) and the 1240K dataset for downstream analyses. We determined genetic sex of each individual by comparing coverage of sex chromosomes with autosomes calculated for the 1240K sites using samtools depth (Figure S1). We detected genetic outliers by projecting Ulsan Koreans to the top principal components calculated for 2,077 present-day Eurasian individuals using the smartpca program v16000 in the EIGENSOFT package v7.2.1 (Patterson et al., 2006). For the remaining individuals, we detected close relative pairs by calculating pairwise mismatch rate (PMR) for each pair using the random haploid calls and by estimating kinship coefficients using the --Z-genome module in PLINK v1.90b6.9 (Chang et al., 2015). We removed one individual from each relative pair up to the second-degree relatives for the downstream group-based analyses (Figure S2; Table S1).

### Data set compilation

We merged genome-wide genotype data of 2,967 present-day individuals typed on the HumanOrigins array (Jeong et al., 2019; Lazaridis et al., 2016; Patterson et al., 2012) with whole genome sequences of 104 Koreans from Ulsan (J. Kim et al., 2018), six Gunsan jar coffin individuals from this study, and previously published ancient individuals (Allentoft et al., 2015; Damgaard, Marchi, et al., 2018; Damgaard, Martiniano, et al., 2018; Fu et al., 2014; Fu et al., 2016; Haber et al., 2017; Harney et al., 2018; Jeong et al., 2016; Jeong et al., 2020; Jeong et al., 2018; Jones et al., 2015; Kanzawa-Kiriyama et al., 2019; Krzewińska et al., 2018; Lazaridis et al., 2017; Lazaridis et al., 2016; Lazaridis et al., 2014; Lipson et al., 2018; Mathieson et al., 2018; Mathieson et al., 2015; McColl et al., 2018; Moreno-Mayar et al., 2018; Narasimhan et al., 2019; Ning et al., 2020; Raghavan, DeGiorgio, et al., 2014; Raghavan, Skoglund, et al., 2014; Rasmussen et al., 2014; Rasmussen et al., 2010; Rasmussen et al., 2015; Sikora et al., 2019; Unterlander et al., 2017; C.-C. Wang et al., 2020; T. Wang et al., 2021; M. A. Yang et al., 2020; Melinda A Yang et al., 2017; Yu et al., 2020). We also merged present-day world-wide populations genotyped on the 1240K sites (Mallick et al., 2016) with whole genome sequences of 104 Koreans from Ulsan, six Gunsan jar coffin individuals from this study, and previously published ancient individuals to produce 1240K dataset. We provide a list of analysis groups and individuals used for each analysis (Table S2).

### Principal component analysis

We ran principal component analysis (PCA) with 2,077 present-day Eurasian individuals from HumanOrigins dataset with the option “lsqproject: YES” using smartpca v16000 and projected the remaining individuals on to the PCs, including the Gunsan jar coffin individuals, present-day Ulsan Koreans, and other ancient East Asian individuals. For the East Asian-only PCA, we ran PCA with 455 present-day East Asian individuals including Ulsan Koreans with the option “lsqproject: YES” and “shrinkmode: YES” using smartpca v16000 and projected the remaining individuals onto the PCs.

### Genetic kinship analysis

We calculated PMR between each pair of ancient Gunsan jar coffin individuals across the 1,150,639 autosomal SNPs in the 1240K panel. Each pair was covered by at least 66,442 SNPs. We estimated probability of sharing 0, 1, and 2 alleles using lcMLkin v0.5.0 to distinguish between parent-offspring and full sibling (Lipatov et al., 2015). For the group-based analyses, we removed first-degree relatives and kept three individuals (GUC002, GUC003, GUC005) with minimal genetic relatedness (Table S3)

### Runs of Homozygosity analysis

We investigated the runs of homozygosity (ROH) segments within the genome of each Gunsan jar coffin individual to understand parental relatedness using hapROH (downloaded from https://pypi.org/project/hapROH/v0.3a1) (Ringbauer et al., 2020). We analyze individuals whose SNPs covered over 400,000 sites among 1240K sites at least once as recommended.

### F-statistics and qpWave/qpAdm analysis

We obtained *f*-statistics using 1240K dataset to maximize SNP coverage of ancient individuals. We calculated outgroup-*f_3_* statistics of the form *f_3_*(Mbuti; Gunsan jar coffin, world-wide) using qp3Pop v650 to measure genetic affinity between two populations. We calculated *f_4_* statistics of the form *f_4_*(Mbuti, world-wide; Gunsan jar coffin, present-day Korean) using qpDstat v970 with the option “f4mode: YES”. We used 121 present-day world-wide populations and 166 ancient populations for these calculations (Table S2). We tested various admixture models of ancient and present-day East Asian populations using qpWave v1200 and qpAdm v1201 programs in the AdmixTools v7.0 (Lazaridis et al., 2016; Reich et al., 2012) on 1240K dataset. We used ten populations as an outgroup set (“right” populations): central African Mbuti (Mbuti.DG; n=5), Early Neolithic farmers from western Anatolia (Anatolia_N; n=23), Andamanese islanders Onge (Onge.DG; n=2), Neolithic Iranians from the Ganj Dareh site (Iran_N; n=8), Epi-Paleolithic European hunter-gatherer from the Villabruna site (Villabruna; n=1), a Late Pleistocene Native American individual from the Upward Sun River site in Alaska (USR1; n=1), Early Neolithic hunter-gatherers from the western Baikal region (Baikal_EN; n=18), an Early Neolithic individual from Shandong region in China (Boshan; n=1), an Early Neolithic individual from the southern Chinese Liangdao site (Liangdao2; n=1), and Funadomari Jomon (Jomon_Funadomari; n=2). We included “allsnps: NO” option.

## Results

### Archaeological context of the Gunsan jar coffin

The Dangbuk-ri site is located in Gunsan, Jeollabuk-do province on the Korean peninsula (Figure 1). The site was salvage excavated for the construction of a railroad line in 2016 by the Central Institute of Cultural Heritage. The archaeological investigation was conducted during the period of approximately 60 days, starting from July 2016. Geographically the site was on the southern hillside of Mt. Dottae, and it covered the top of a hill, the height of which was about 60 meters above sea level. A total of 21 burials were found at the Dangbuk-ri site. Among those burials, sixteen were the stone-cist type, four burials were the stone-chamber type, and the remaining one was a jar coffin. Archaeological contexts, such as burial types and artifacts, suggest that these burials are dated to the Woongjin period (475-538 AD) and the Sabi period (538-660 AD) of Baekje, corresponding to the early Medieval Three Kingdoms Period.

The jar coffin burial was located on the southeastern hillside from the top, surrounded by stone-cist burials and stone-chamber burials without any specific pattern. The jar was laid in a circular pit and two stones were placed beside the jar to fix the jar body. At the time of discovery, the jar was found slanted slightly. The mouth area of the jar has been broken without a cover of the coffin. In the jar coffin, multiple skeletons were found together (Figure 2). No specific pattern was detected in the placement of skeletal elements within the jar coffin, although more skulls were placed in the upper part of the jar. No case of a sequential multiple burial (i.e. individuals were sequentially put to the jar coffin over an extended time period) has been reported for this type of jar coffins, although there are cases with those consisting of two jars. Also, individuals in this jar coffin were not separated into layers, a pattern expected for a sequential multiple burial. Thus, multiple skeletons in the jar coffin are considered as interred together approximately at the same time as a secondary burial. After finishing the excavation, a pottery as a grave good was found at the bottom of a circular pit, implying that ritual behavior was performed before laying the jar-coffin in state. The height of the jar, from the lower part of the jar to the top is 72.3cm, the diameter is 32.1cm.

### Ancient genome-wide data production

We performed a genetic investigation of early Medieval individuals from Dangbuk -ri, Gunsan, in the southwestern region of the Republic of Korea (Figure 1; Table 1). These individuals are dated to 5^th^ to 7^th^ century AD based on archaeological contexts and are found in a single jar coffin, with signatures of secondary burial: multiple burials, either primary or secondary, within a single jar coffin are exceptional given that most jar coffins host only a single individual. Based on the macroscopic examination of all skeletal elements retrieved from the jar coffin, we determined MNI as nine, including six adults and three subadults (Table 1). Following an in-solution capture protocol for ~1.2 million informative SNPs (Mathieson et al., 2015), we obtained genome-wide data with on-target autosomal coverage ranging 0.2-1.8x and with 189K-764K on-target SNPs covered at least once for six individuals (Table 2). The remaining three of nine individuals in the jar coffin did not yield sufficient genomic DNA to be analyzed. All six individuals show post-mortem chemical damages typical of aDNA. All four males show nuclear contamination < 2% based on their X chromosome data and five of six individuals have mitochondrial contamination of 1-3%. Based on these measures, we included all six individuals into the downstream analysis, including the one female without a contamination measure due to low mitochondrial coverage (Table 2). For each individual, we called haploid genotypes across the 1240K panel SNPs by randomly sampling a high-quality base. We concatenated their genotype data with published presenet-day and ancient individuals on the HumanOrigins and the 1240K data sets for downstream analyses (Table S2)

**Table 1.**
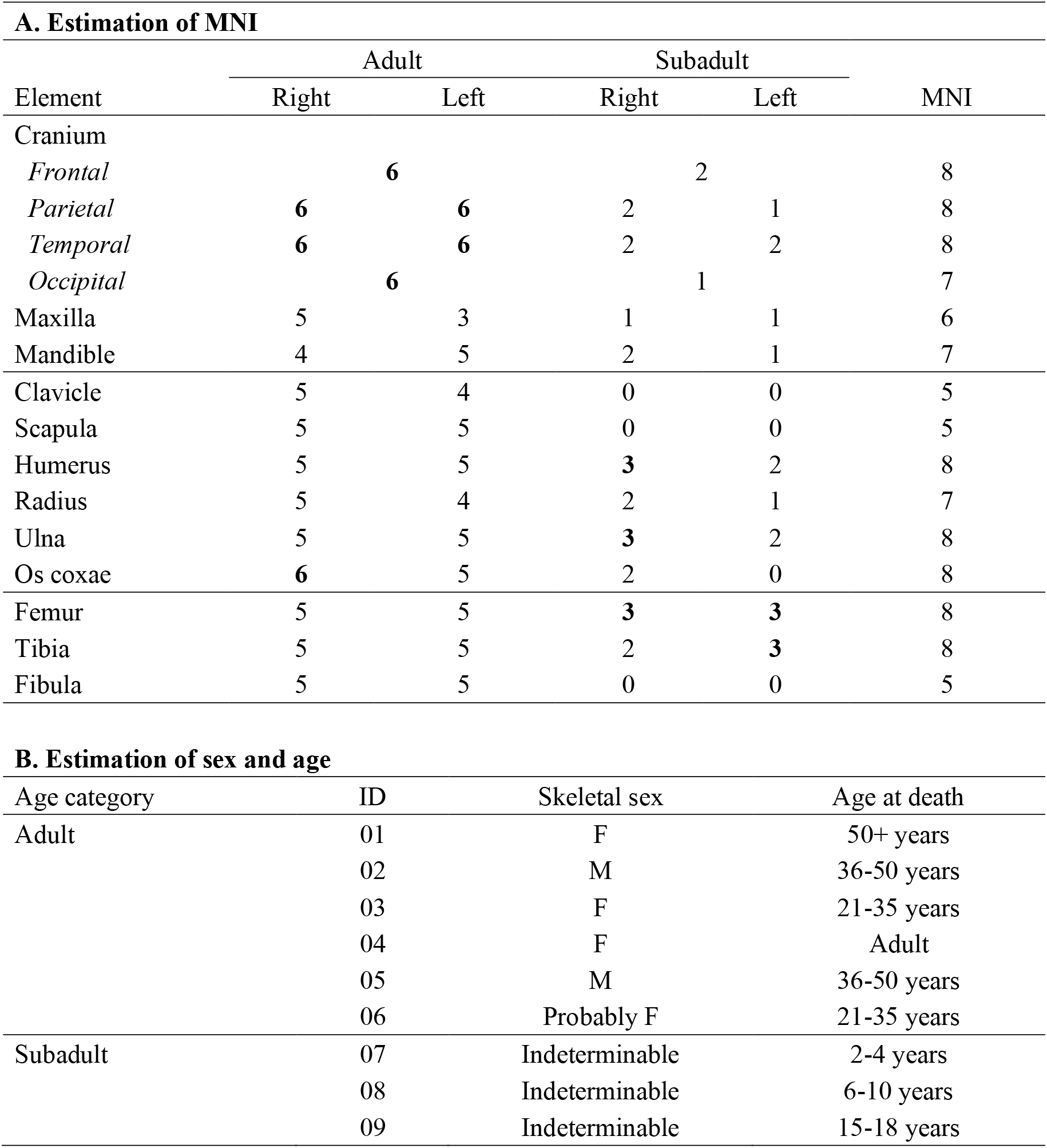
Summary of morphological examination of the skeletal elements found in the Gunsan jar coffin. (A) Estimation of MNI based on the number of skeletal elements. We detected at least six adult individuals and three subadults, totaling at least nine individuals in the jar coffin. Because of incomplete preservation of skeletal elements per individual, per-element MNI is smaller than the MNI based on all elements. (B) Estimation of sex and age of each individual.

**Table 2.** A summary of sequencing and genetic information of six ancient individuals in this study. Six of the seven individuals yield sufficient human DNA for genome-wide analysis. The numbers of covered target sites are counted among 1,233,013 SNPs in the 1240K panel and 593,124 autosomal SNPs in the HumanOrigins panel. X-chromosome based contamination estimates represent the point estimate ± 1 s.e.m., and mitochondrial estimates represent the point estimate and the 95% credible interval.

| Lab ID | ID | Genetic Sex | Pre-capture % human DNA | Post-capture % target reads | # of reads sequenced | # of uniquely mapped reads |
| --- | --- | --- | --- | --- | --- | --- |
| GUC001 | 01; skull 1 | M | 0.129 | 2.45 | 36,063,678 | 1,169,573 |
| GUC002 | 02; skull 2 | M | 1.219 | 14.90 | 25,871,919 | 5,363,973 |
| GUC003 | 03; skull 3 | M | 0.307 | 4.99 | 32,502,379 | 2,076,521 |
| GUC004 | 04; skull 4 | F | 0.350 | 6.89 | 36,179,211 | 2,592,575 |
| GUC005 | 06; skull 6 | M | 2.624 | 22.91 | 20,992,582 | 6,805,731 |
| GUC007 | 09; subadult 3 | F | 1.656 | 18.35 | 24,249,836 | 5,438,347 |
| GUC006 | 08; subadult 2 | N/A | 0.030 | Fail | N/A | N/A |

| Lab ID | Lat | Lon | Coverage |  |  |  | # of covered target sites |  |
| --- | --- | --- | --- | --- | --- | --- | --- | --- |
|  |  |  | Autosome | X | Y | MT | 1240K | HumanOrigins |
| GUC001 | 35.94038889 | 126.7141944 | 0.195 | 0.075 | 0.091 | 0.54 | 189,920 | 96,738 |
| GUC002 | 35.94038889 | 126.7141944 | 1.097 | 0.420 | 0.518 | 3.11 | 639,758 | 327,372 |
| GUC003 | 35.94038889 | 126.7141944 | 0.312 | 0.122 | 0.147 | 0.48 | 284,857 | 144,896 |
| GUC004 | 35.94038889 | 126.7141944 | 0.397 | 0.304 | 0.004 | 0.56 | 342,704 | 174,677 |
| GUC005 | 35.94038889 | 126.7141944 | 1.839 | 0.698 | 0.939 | 2.17 | 764,537 | 389,635 |
| GUC007 | 35.94038889 | 126.7141944 | 1.087 | 0.812 | 0.015 | 2.22 | 643,530 | 330,307 |

| Lab ID | Post-mortem damage |  | Contamination estimates |  | Uniparental haplogroup |  |
| --- | --- | --- | --- | --- | --- | --- |
|  | 5' C>T | 3' G>A | X | MT | MT | Y |
| GUC001 | 0.2928 | 0.2819 | 0.0184±0.0147 | 0.01 (0.00-0.02) | D4c1b1 | Q1a (Q-L472) |
| GUC002 | 0.2096 | 0.1952 | 0.0020±0.0017 | 0.01 (0.00-0.02) | D4b2b1 | Q1a1a1 (Q-M120) |
| GUC003 | 0.2623 | 0.2575 | 0.0106±0.0073 | 0.01 (0.00-0.02) | B5b3a | Q1a1 (Q-F1251) |
| GUC004 | 0.2413 | 0.2376 | N/A | N/A | N/A | - |
| GUC005 | 0.2140 | 0.2012 | 0.0058±0.0016 | 0.03 (0.01-0.05) | D4c1b1 | O1b2a2a1a (O-CTS7620) |
| GUC007 | 0.2328 | 0.2119 | N/A | 0.02 (0.00-0.04) | D4c1b1 | - |

### A familial relationship of individuals from a single jar coffin

We first ran PCA of 2,077 present-day Eurasians and projected ancient individuals from the Gunsan jar coffin onto the top PCs (Figure 3). All six individuals fall close to each other and to present-day Koreans, suggesting no substantial genetic heterogeneity among them and an overall close relationship with present-day Koreans (Figure 3). In the PCA of present-day East Asians including present-day Koreans, the Gunsan jar coffin individuals also overlap with present-day Koreans (Figure S3).

**Figure 3.**
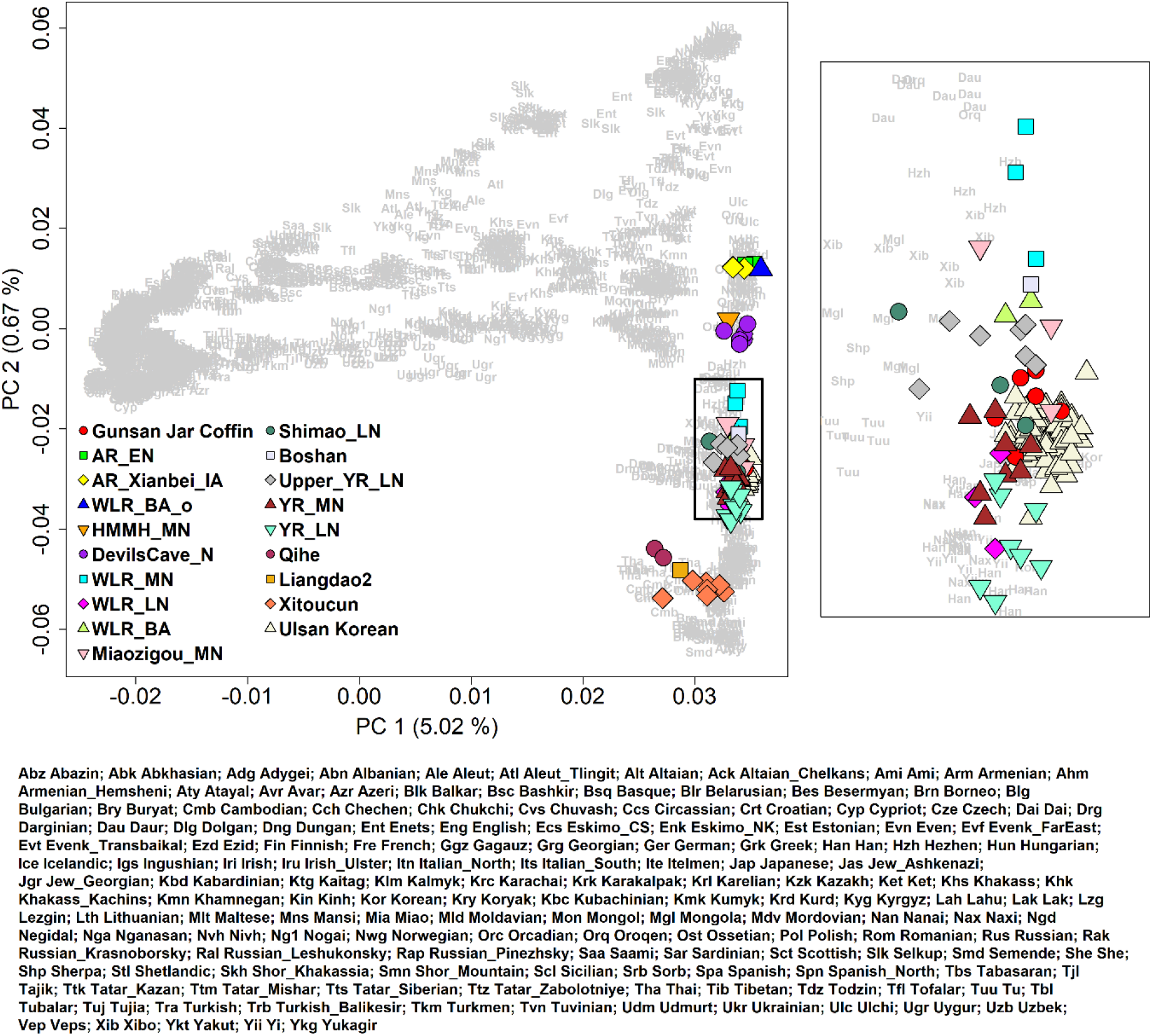
Principal component analysis from 2,077 present-day Eurasian individuals. We project the Gunsan jar coffin and other ancient East Asian individuals and present-day Koreans from Ulsan (marked by color-filled shapes) onto the top two PCs calculated for 2,077 present-day Eurasian individuals (marked by three-letter codes). Present-day and ancient Koreans fall on top of each other.

Then we measured the genetic relatedness between these ancient individuals to test if the unusual occasion of multiple individuals in a single jar coffin reflects their close relationship. Our estimates of genetic relatedness based on genome-wide data indeed show that these individuals form a closely related extended family: among 15 pairs combined from six individuals, we observe six first-degree, two second-degree, and three third-degree relative pairs (Figure 4A; Table S3). Incorporating mitochondrial and Y haplogroup information and age at death, we distinguish between parent-offspring and full sibling pairs and propose most plausible pedigrees (Figure 4B). At the core of this pedigree is a quartet family composed of a couple and their two children, one adult male and one subadult female. Among the remaining two individuals, one male (GUC005) is the 1^st^ degree relative of the mother of the quartet (GUC004) and the 2^nd^ degree relative of the two children (GUC001 and GUC007), but is unrelated to the father (GUC002). The 1^st^ degree relationship of GUC005 and GUC004 is likely a parent-offspring one given the near-zero sharing of both alleles at the same SNP (Table S3). Both father-daughter and mother-son relationship are compatible with genetic data: however, given that GUC005 and the quartet children share the identical mitochondrial haplogroup, we propose mother-son may be more likely. That is, GUC005 and (GUC001, GUC007) may be half-siblings with the same mother. Lastly, GUC003 is most likely a 3^rd^ degree relative of the quartet father, sharing the same Y haplogroup.

**Figure 4.**
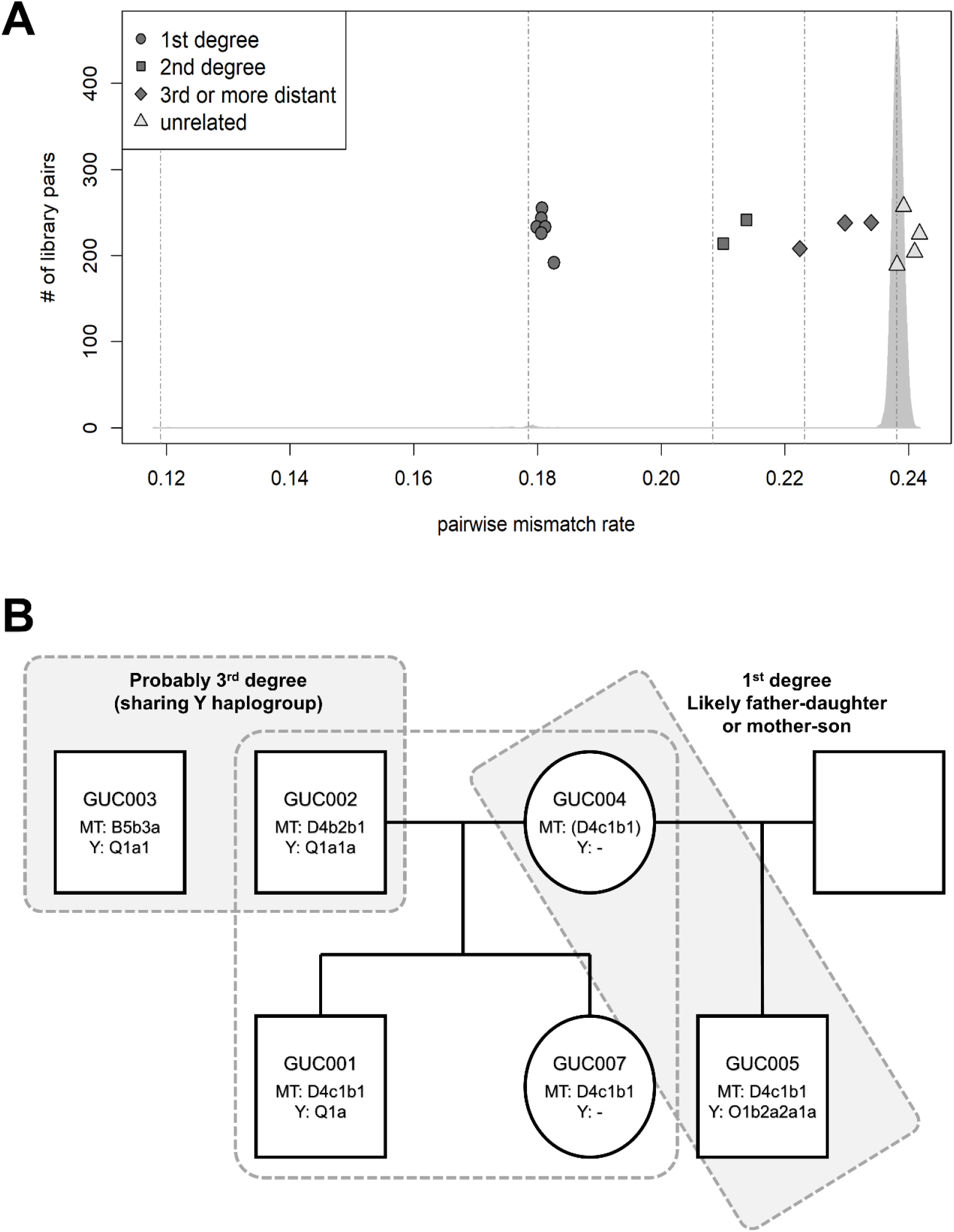
Genetic relationship of the six Gunsan jar coffin individuals. (A) We show the estimated genetic relatedness of 15 pairs of the Gunsan jar coffin individuals with the pairwise mismatch rate of genotypes (color-filled shapes). On the background, we plot the density of the pairwise mismatch rate values of 102 present-day Ulsan Koreans. Dotted vertical lines represent the expected pairwise mismatch rate of the identical, 1^st^ degree, 2^nd^ degree, 3^rd^ degree relatives and the unrelated pairs of present-day Koreans, from left to right, respectively. Gunsan jar coffin individuals show slightly higher pairwise mismatch rate values than the present-day Koreans. (B) A reconstruction of the pedigree of the six Gunsan jar coffin individuals. MT and Y represent the corresponding uniparental haplogroups. GUC004 and GUC005 are the 1^st^ degree relatives that are likely mother-son or father-daughter. Here we show a mother-son relationship based on the shared MT haplotype between GUC005 and (GUC001, GUC007), offspring of GUC004.

We also investigated the distribution of ROH segments in three individuals with sufficient coverage (> 400K of 1240K SNPs are covered) using hapROH (Ringbauer et al., 2020). Finding few long ROH segments (> 4 cM), we conclude that none of these individuals were the offspring of close relatives.

### The genetic profile of early Medieval Koreans

To understand the genetic profile of the early Medieval Koreans from the Gunsan jar coffin, we first measured the genetic affinity between the Gunsan jar coffin group and world-wide present-day and ancient populations using outgroup-*f_3_* statistics of the form *f_3_*(Mbuti; Gunsan jar coffin, world-wide). As expected, the Gunsan jar coffin group shows the highest genetic affinity with present-day Koreans (Figure. S4). Formally testing the genetic symmetry of the early Medieval and present-day Koreans using *f_4_* statistics of the form *f_4_*(Mbuti, world-wide; Gunsan jar coffin, present-day Korean), only a few groups break the symmetry without clear geographic distribution (Figure S5). To formally model the genetic relationship between ancient and present-day Korean groups, we utilized qpAdm, which summarizes multiple *f_4_* statistics to test if a mixture of allele frequency of the chosen source populations can accurately mimic that of the target population (Lazaridis et al., 2016; Reich et al., 2012). We find that the present-day Koreans from Ulsan city (n=88) are adequately modeled as a mixture of Gunsan jar coffin group and a European source with a small negative coefficient (−1.0% to −1.4%; Table S4A). Reciprocally, Gunsan jar coffin group is modeled as a mixture of present-day Ulsan Koreans and a small contribution from a European source (1.0-1.4%; Table S4A). We interpret these results as a technical artifact rather than a signal of true admixture, considering a comparable amount of nuclear contamination in Gunsan individuals (0.2-1.8%) and the small reference bias introduced during the read mapping step.

To compare the genetic profiles of the ancient and present-day Koreans with populations of the surrounding regions in a broader sense, we searched for distal admixture models that were commonly applicable to those populations. We find that the Gunsan jar coffin individuals, present-day Koreans, and various ancient groups from northern China are adequately positioned along the genetic north-south cline in East Asia, within the range defined by the following two populations: i) Bronze Age individuals from the Longtoushan archaeological site of the West Liao River region in the Upper Xiajiadian culture context (“WLR_BA”) as a genetic northern proxy, and ii) Late Neolithic individuals from the Xitoucun site in southern China (“Xitoucun”) as a genetic southern proxy (Figure 5; Table S5A). In both ancient and present-day Koreans, we do not detect a statistically significant contribution from the Jomon hunter-gatherer gene pool of the Japanese archipelago (Table S5A), although previous studies report occasional presence of the Jomon ancestry contribution from Neolithic to the early Medieval period (Gelabert et al., 2021; Robbeets et al., 2021). When we replace the genetic northern proxy from WLR_BA to Middle Neolithic individuals from the Miaogizou site in Inner Mongolia (“Miaozigou_MN”), we detect a small but significant amount of Jomon contribution in the Gunsan individuals and present-day Ulsan Koreans (3.1-4.4%; Table S5B). We believe that WLR_BA provides a more suitable model for ancient and present-day Koreans given its geographical and temporal proximity to them.

**Figure 5.**
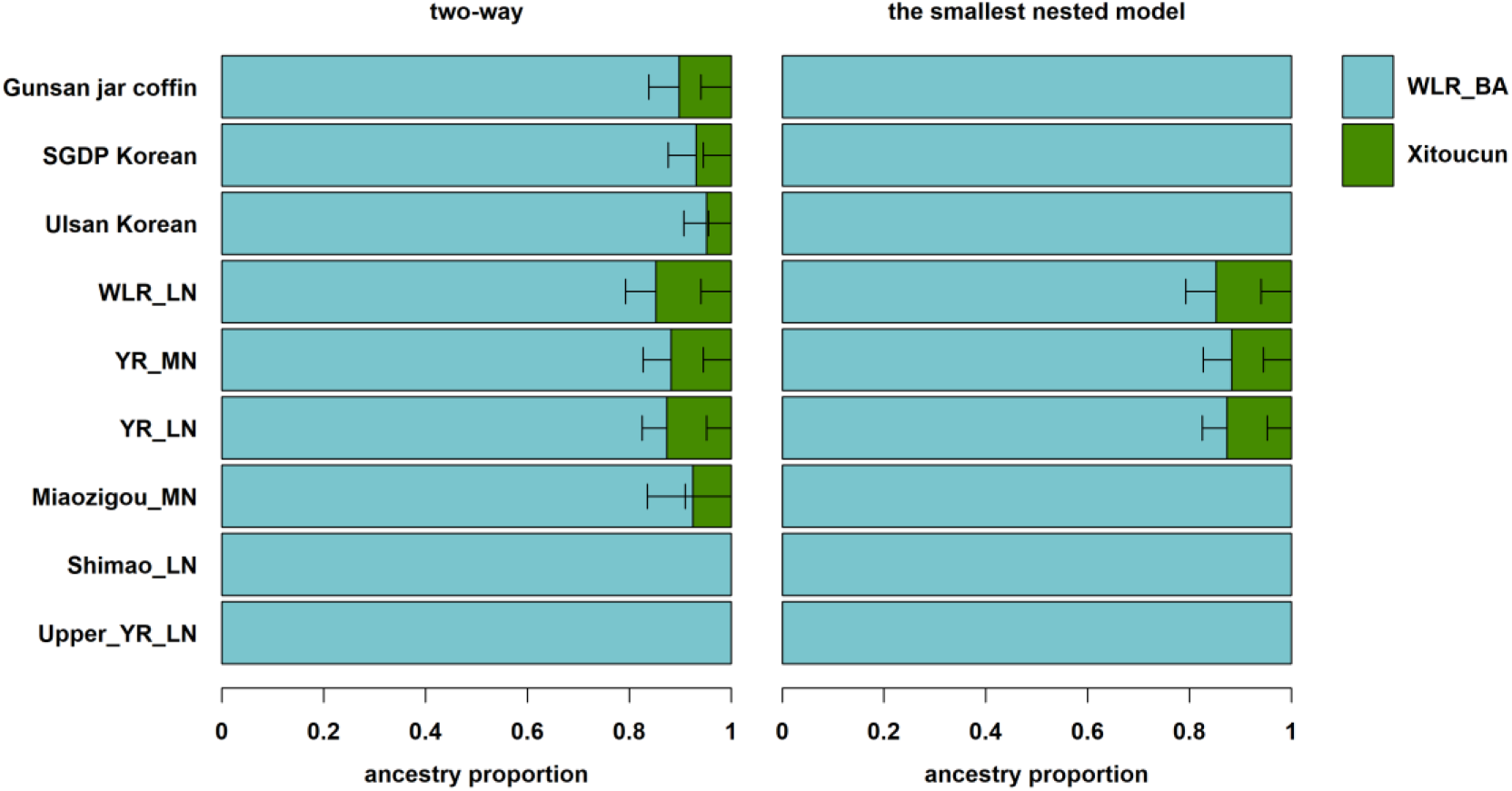
QpAdm modeling of Gunsan jar coffin and other ancient and present-day East Asian populations. We show the results of a two-way admixture model of WLR_BA+Xitoucun on the left side and the results of its smallest sub-model after removing all components that do not significantly increase model fit on the right side (Table S5A). Horizontal bars represent the standard error measure (s.e.m.) of ancestry proportion estimates, calculated by the 5cM block jackknifing procedure as implemented in the qpAdm program.

## Discussion

We present a genomic study of individuals from the Gunsan jar coffin, an unusual case of a secondary multiple burial where at least nine individuals were co-buried in a single small jar coffin. We confirm our hypothesis that this unusual burial represents an unusual relationship among the co-buried individuals, i.e. an extended family, including a core of a couple and their two children, as well as both paternal and maternal relatives. The inferred pedigree may imply little gender bias in the family structure of early Medieval Koreans lived in the area. Further archaeogenomic studies on the ancient individuals excavated from an unusual burial context or those from a single cemetery will provide more insights into past mortuary practices and social structures.

While the genetic origins of present-day Koreans have not been fully understood due to lack of relevant ancient genome data, individuals from the Gunsan jar coffin provide among the first past human genetic profiles in ancient Korea. Since this type of small jar coffins are considered to be associated with commoners rather than with sociopolitical elites, the Gunsan jar coffin individuals provide a glimpse on the genetic profile of the general population of the early Medieval Korea, albeit small in number. Our population genomic analysis shows a long-term presence of the genetic profile of present-day Koreans in the Korean peninsula at least since the early Medieval period. However, this does not imply a complete genetic isolation of the Korean population from their neighbors over the last two millennia. On the contrary, there are a growing body of genetic evidence supporting high connectivity between proto-historic Korea and its neighboring regions: a recent study reported a few early Medieval individuals with a substantial level of the Jomon ancestry from the Japanese archipelago (Gelabert et al., 2021), suggesting a vibrant international network supporting movements of people and goods. Furthermore, Kofun-period individuals from Japan suggests a continued gene flow from the continental East Asia with the Korean peninsula as a highly likely source region (Cooke et al., 2021). Further archaeogenetic studies on proto-historic sites in and around the Korean peninsula will help us accurately delineate the networks between the past East Asian societies.

## Acknowledgements

This work was supported by the National Research Foundation of Korea grant funded by the Korea government (No. 2018R1A5A7023490 to E.J.W. and 2020R1C1C1003879 to C.J.) and the Max Planck Society.

## Data Availability

The raw DNA sequences (FASTQ) and the alignment data (BAM) reported in this paper have been deposited in the European Nucleotide Archive under the accession number PRJEB51247. Data will be made publicly available when the manuscript is published.

## Conflict of Interest Statement

The authors declare no conflict of interest.

## Notes

### Competing Interest Statement

The authors have declared no competing interest.

