## Supplementary material for "Genomic detection of a secondary family burial in a single jar coffin in early Medieval Korea": Figure S1, Figure S2, Figure S3, Figure S4, Figure S5, Table S1, Table S3, Table S4, Table S5

**This file includes:**

Figures S1 to S5
Tables S1 to S5

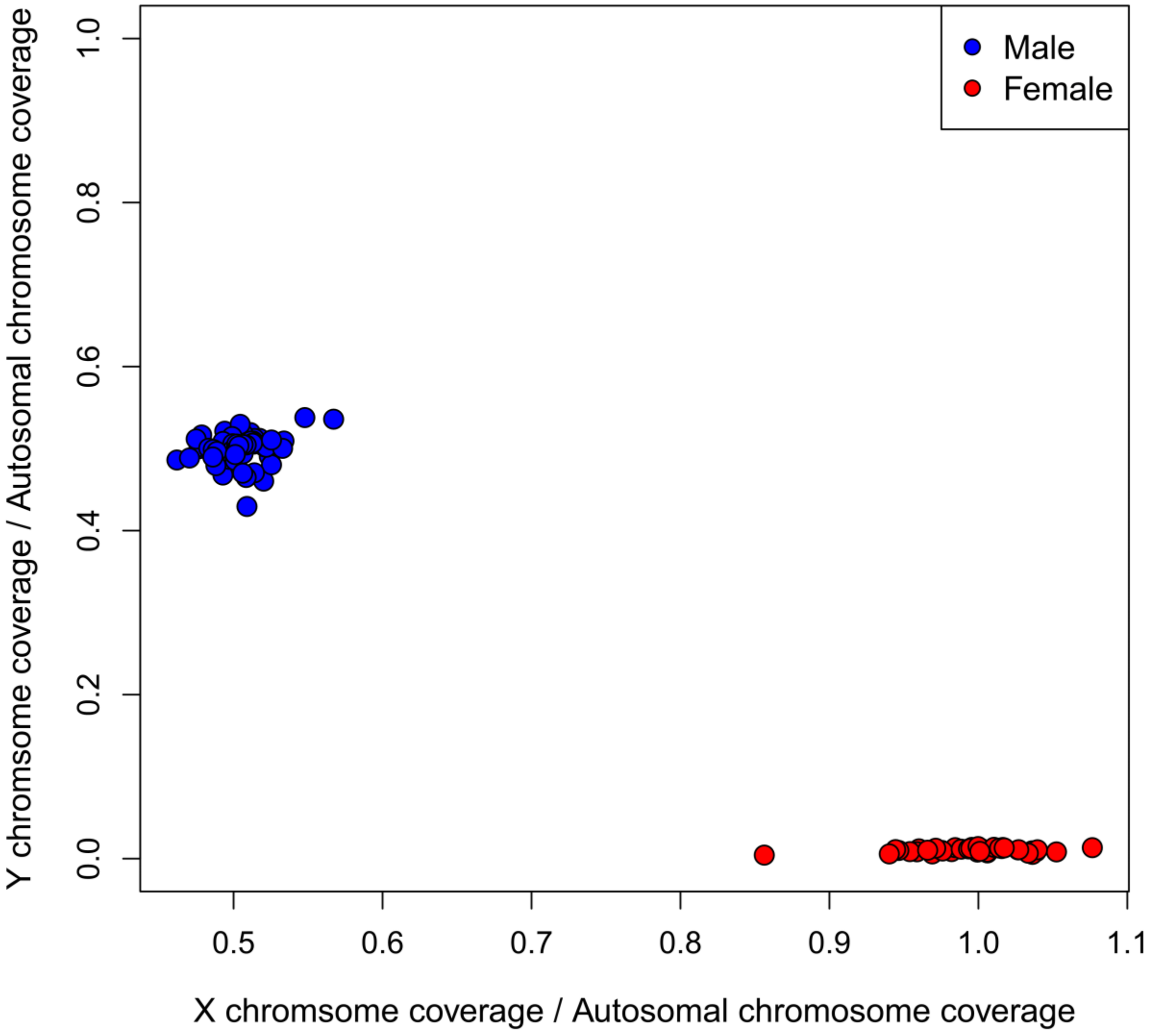


**Figure S1. Genetic sex assignment of present-day Koreans from Ulsan.** We plot the ratio of X to autosomal coverage (x-axis) and Y to autosomal coverage (y-axis). Blue and red circles represent genetic males (XY) and females (XX), respectively. No individual with obvious sex chromosome aneuploidy is observed.

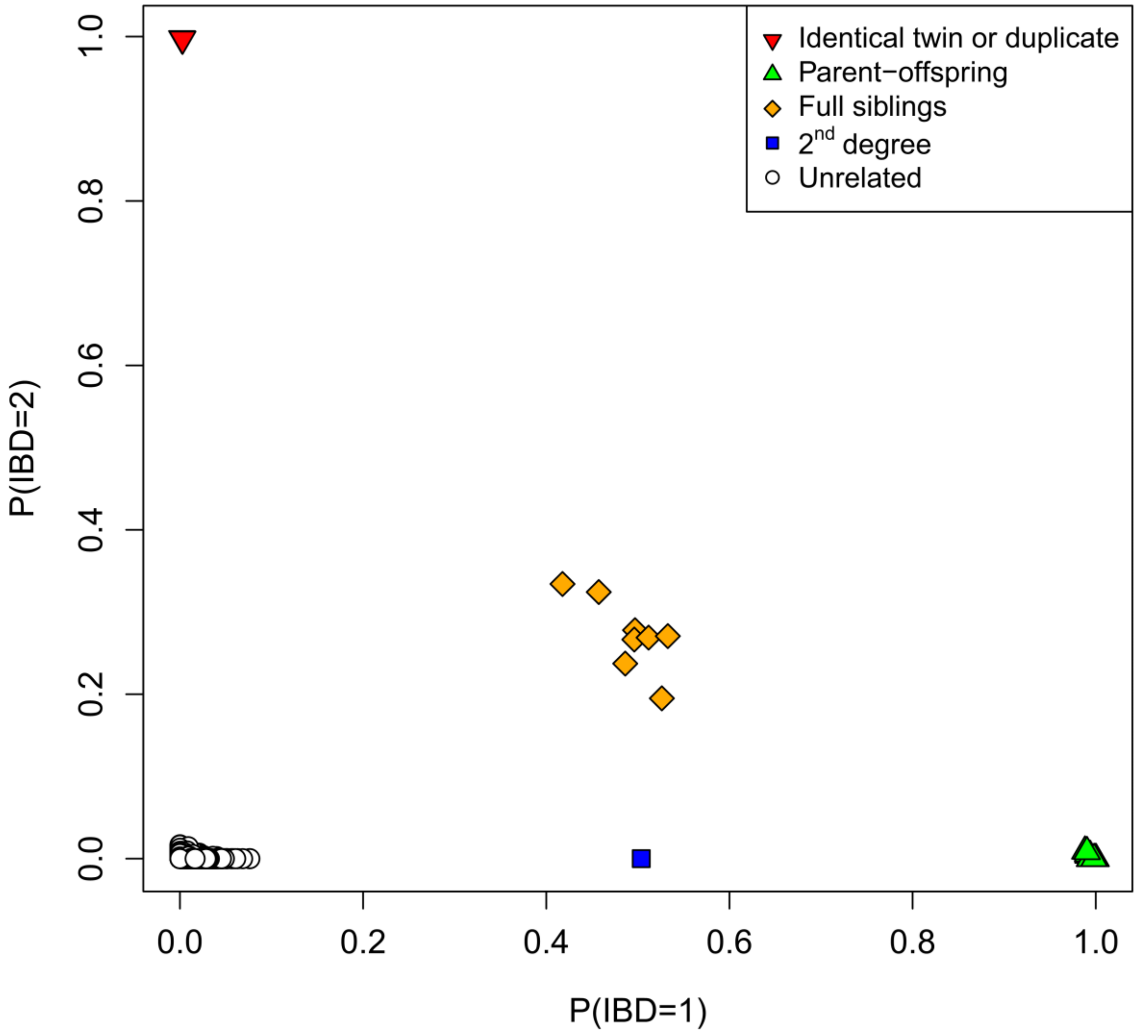


**Figure S2. Classification of relatives among present-day Ulsan Korean individuals.** We plot PLINK estimates of the probability of sharing one and two alleles per each SNP.

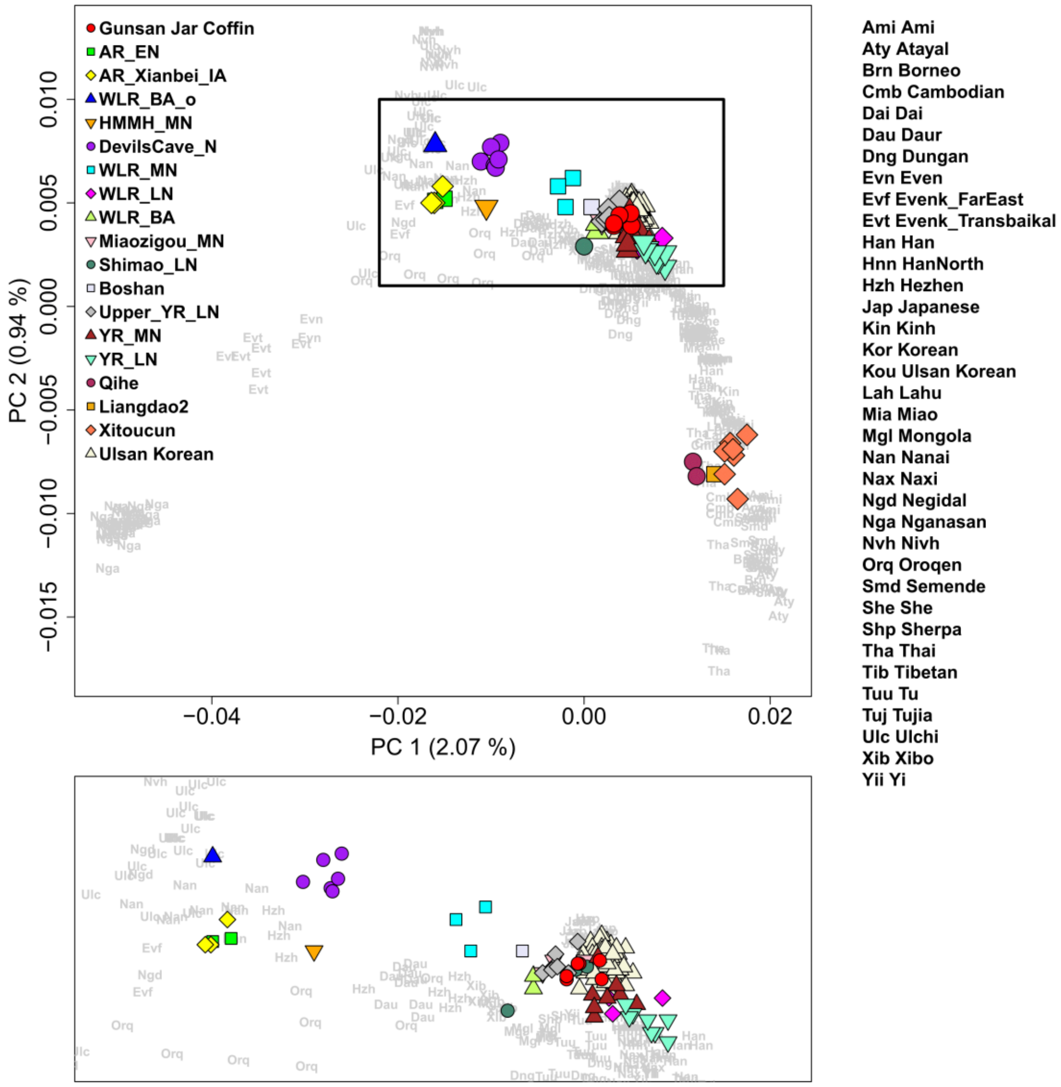


**Figure S3. Principal component analysis from 455 present-day East Asian individuals.** We project the Gunsan jar coffin and other ancient East Asian individuals (marked by color-filled shapes) onto the top two PCs calculated for 455 present-day East Asian individuals (marked by three-letter codes). Ulsan Koreans are also marked by triangle shapes. Present-day and ancient Koreans fall on top of each other.

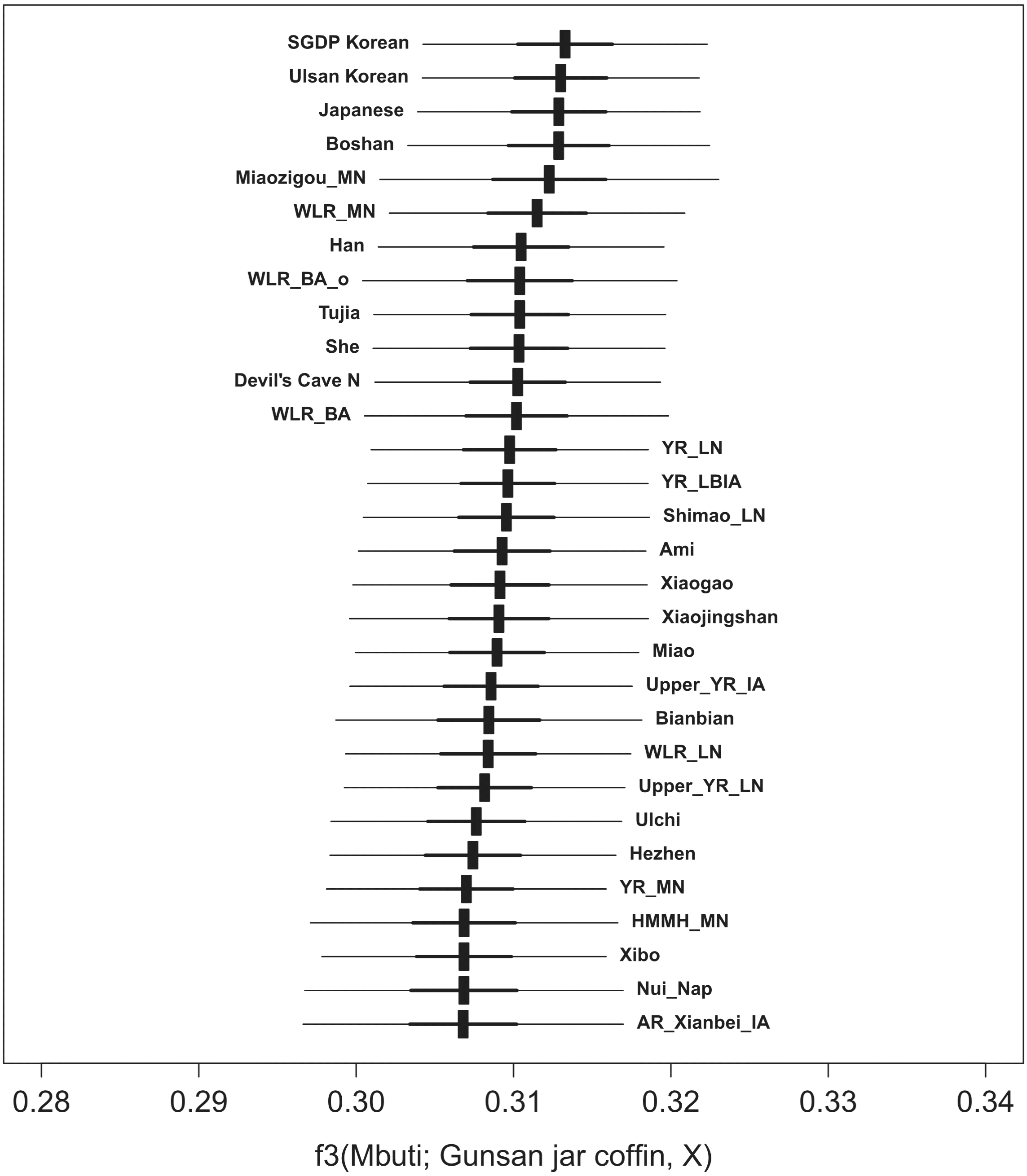


**Figure S4. Top 30 outgroup-$f_3$ statistics of the form $f_3$(Mbuti; Gunsan jar coffin, world-wide) for ancient and modern worldwide populations.** Gunsan jar coffin individuals show the highest genetic affinity with present-day Koreans, followed by other ancient and present-day East Asians. Horizontal bars represent the point estimate ± 3 (thin) and ± 1 (thick) standard error measure (s.e.m.), respectively. s.e.m. are calculated by 5cM block jackknifing.

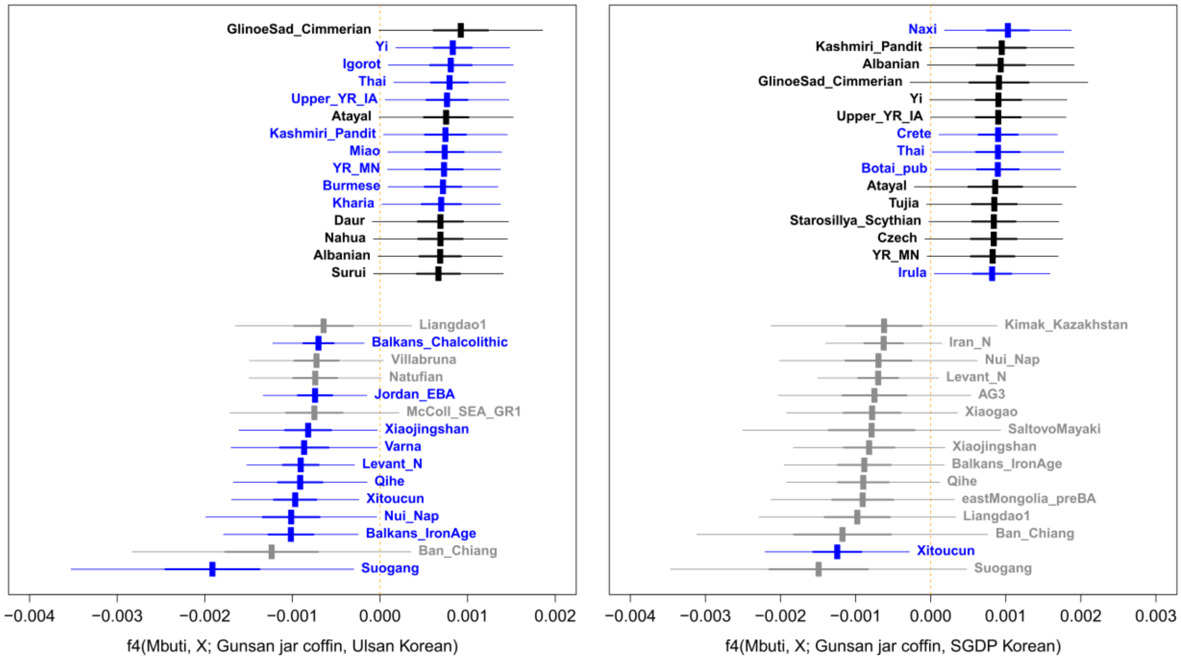


**Figure S5. A comparison of the genetic affinity of early Medieval and present-day Koreans with world-wide populatios.** We present the 15 most positive and negative $f_4$ statistics of the form $f_4$(Mbuti, X; Gunsan jar coffin, present-day Korean). $F_4$ statistics with |Z| > 3 are marked by blue color. Horizontal bars represent the point estimate ± 3 (thin) and ± 1 (thick) standard error measure (s.e.m.), respectively. s.e.m. are calculated by 5cM block jackknifing.

**Table S1. Genetic relatives detected among the 104 present-day Koreans from Ulsan.** We show all relative pairs up to the 2nd degree relatives. Z0, Z1, Z2 represent PLINK estimates of the probability of sharing 0, 1, 2 alleles, respectively. PI_HAT represents the PLINK estimate of the genetic relatedness. PMR represents the pairwise mismatch rate of genotypes.

| ID1 | ID2 | Z0 | Z1 | Z2 | PI_HAT | PMR | Kinship |
|---|---|---|---|---|---|---|---|
| 00088 | 00089 | 0.0001 | 0.0022 | 0.9976 | 0.9988 | 0.1197 | identical |
| 00090 | 00091 | 0.0002 | 0.0028 | 0.9970 | 0.9984 | 0.1216 | identical |
| 00231 | 00252 | 0.2183 | 0.4573 | 0.3243 | 0.5530 | 0.1728 | full siblings |
| 00252 | 00253 | 0.2482 | 0.4177 | 0.3341 | 0.5430 | 0.1742 | full siblings |
| 00337 | 00338 | 0.1966 | 0.5327 | 0.2707 | 0.5370 | 0.1749 | full siblings |
| 00003 | 00009 | 0.2253 | 0.4969 | 0.2779 | 0.5263 | 0.1767 | full siblings |
| 00219 | 00221 | 0.2194 | 0.5117 | 0.2689 | 0.5247 | 0.1772 | full siblings |
| 00005 | 00009 | 0.2374 | 0.4961 | 0.2665 | 0.5145 | 0.1775 | full siblings |
| 00231 | 00253 | 0.2765 | 0.4862 | 0.2373 | 0.4804 | 0.1819 | full siblings |
| 00003 | 00005 | 0.2789 | 0.5262 | 0.1950 | 0.4581 | 0.1843 | full siblings |
| 00353 | 00362 | 0.0012 | 0.9904 | 0.0084 | 0.5036 | 0.1786 | parent-child |
| 00002 | 00005 | 0.0023 | 0.9938 | 0.0039 | 0.5008 | 0.1787 | parent-child |
| 00337 | 00343 | 0.0015 | 0.9985 | 0.0000 | 0.4993 | 0.1788 | parent-child |
| 00001 | 00003 | 0.0044 | 0.9956 | 0.0000 | 0.4978 | 0.1795 | parent-child |
| 00342 | 00343 | 0.0019 | 0.9981 | 0.0000 | 0.4991 | 0.1795 | parent-child |
| 00232 | 00234 | 0.0028 | 0.9882 | 0.0090 | 0.5031 | 0.1796 | parent-child |
| 00002 | 00009 | 0.0000 | 1.0000 | 0.0000 | 0.5000 | 0.1797 | parent-child |
| 00001 | 00009 | 0.0049 | 0.9951 | 0.0000 | 0.4976 | 0.1798 | parent-child |
| 00228 | 00253 | 0.0067 | 0.9933 | 0.0000 | 0.4966 | 0.1799 | parent-child |
| 00234 | 00235 | 0.0000 | 1.0000 | 0.0000 | 0.5000 | 0.1799 | parent-child |
| 00230 | 00231 | 0.0032 | 0.9912 | 0.0056 | 0.5012 | 0.1800 | parent-child |
| 00205 | 00221 | 0.0000 | 1.0000 | 0.0000 | 0.5000 | 0.1800 | parent-child |
| 00002 | 00003 | 0.0000 | 1.0000 | 0.0000 | 0.5000 | 0.1800 | parent-child |
| 00228 | 00252 | 0.0000 | 1.0000 | 0.0000 | 0.5000 | 0.1800 | parent-child |
| 00230 | 00252 | 0.0065 | 0.9935 | 0.0000 | 0.4967 | 0.1800 | parent-child |
| 00228 | 00231 | 0.0000 | 1.0000 | 0.0000 | 0.5000 | 0.1802 | parent-child |
| 00001 | 00005 | 0.0000 | 1.0000 | 0.0000 | 0.5000 | 0.1803 | parent-child |
| 00205 | 00219 | 0.0000 | 1.0000 | 0.0000 | 0.5000 | 0.1806 | parent-child |
| 00230 | 00253 | 0.0000 | 1.0000 | 0.0000 | 0.5000 | 0.1810 | parent-child |
| 00220 | 00221 | 0.0044 | 0.9956 | 0.0000 | 0.4978 | 0.1812 | parent-child |
| 00219 | 00220 | 0.0043 | 0.9917 | 0.0040 | 0.4998 | 0.1813 | parent-child |
| 00231 | 00253 | 0.2765 | 0.4862 | 0.2373 | 0.4804 | 0.1819 | full siblings |
| 00003 | 00005 | 0.2789 | 0.5262 | 0.1950 | 0.4581 | 0.1843 | full siblings |
| 00338 | 00343 | 0.4963 | 0.5037 | 0.0000 | 0.2518 | 0.2081 | 2nd degree |

**Table S2. A list of world-wide ancient and present-day populations used in this study.**
[Please see the excel file]

**Table S3. Genetic relatedness of six Gunsan jar coffin individuals.** We detect six $1^{st}$ degree (five parent-offsprint and one full sibling), two $2^{nd}$ degree, three $3^{rd}$ degree or more distant relative pairs among 15 pairs. We infer genetic relatedness based on the pairwise mismatch rate, and distinguish between parent-offspring and full sibling based on the probability of sharing both alleles estimated by the lcMLkin program using genotype likelihood data. k0, k1, k2 represent probability of sharing 0, 1, and 2 alleles, respectively.

| | | Pairwise Mismatch Rate | | | lcMLkin | | | |
|---|---|---|---|---|---|---|---|---|
| ID1 | ID2 | Covered | Mismatch | Pr(mismatch) | k0 | k1 | k2 | Kinship |
| GUC001 | GUC002 | 135,698 | 24,525 | 0.1807 | 0.293 | 0.640 | 0.067 | Son-Father |
| GUC001 | GUC003 | 66,442 | 15,546 | 0.2340 | 0.992 | 0.008 | 0.000 | > $3^{rd}$ degree |
| GUC001 | GUC004 | 77,168 | 14,096 | 0.1827 | 0.324 | 0.620 | 0.055 | Son-Mother |
| GUC001 | GUC005 | 154,884 | 33,116 | 0.2138 | 0.736 | 0.262 | 0.002 | $2^{nd}$ degree |
| GUC001 | GUC007 | 135,518 | 24,482 | 0.1807 | 0.453 | 0.314 | 0.233 | Full sibling |
| GUC002 | GUC003 | 196,448 | 43,698 | 0.2224 | 0.871 | 0.124 | 0.005 | $3^{rd}$ degree |
| GUC002 | GUC004 | 230,345 | 54,847 | 0.2381 | 0.996 | 0.004 | 0.000 | Unrelated |
| GUC002 | GUC005 | 487,550 | 116,613 | 0.2392 | 0.996 | 0.004 | 0.000 | Unrelated |
| GUC002 | GUC007 | 420,238 | 75,589 | 0.1799 | 0.172 | 0.812 | 0.016 | Father-daughter |
| GUC003 | GUC004 | 110,450 | 26,615 | 0.2410 | 0.995 | 0.005 | 0.000 | Unrelated |
| GUC003 | GUC005 | 225,119 | 54,425 | 0.2418 | 0.995 | 0.004 | 0.000 | Unrelated |
| GUC003 | GUC007 | 196,314 | 45,100 | 0.2297 | 0.978 | 0.022 | 0.000 | > $3^{rd}$ degree |
| GUC004 | GUC005 | 265,920 | 48,192 | 0.1812 | 0.217 | 0.758 | 0.024 | $1^{st}$ degree; likely mother-son or daughter-father |
| GUC004 | GUC007 | 231,027 | 41,733 | 0.1806 | 0.254 | 0.722 | 0.024 | Mother-daughter |
| GUC005 | GUC007 | 489,161 | 102,749 | 0.2101 | 0.670 | 0.318 | 0.013 | $2^{nd}$ degree |

**Table S4. QpAdm-based admixture modeling of early Medieval and present-day Ulsan Koreans using proximal sources.** (A) The Gunsan jar coffin idnviduals are adequatel modeled as present-day Ulsan Koreans with a small amount of European ancestry, here represented by the early Neolithic Central Europeans ("LBK_EN") and by the Middle-Late Bronze Age Russian Steppe population ("Sintashta_MLBA"). This is most likely an artefact due to a small amount of contamination and the reference bias. The Jomon ancestry does not explain the observed difference between the Gunsan jar coffin and present-day Ulsan Koreans. Alternatively, a mixture of Gunsan jar coffin individuals and the Iron Age individuals from Mogushan site ("AR_Xianbei_IA") also fits the present-day Ulsan Koreans. (B) However, a three-way model of Gunsan jar coffin+AR_Xianbei_IA+European shows a similar level of European ancestry and a non-significant contribution from AR_Xianbei_IA, suggesting the artifactual European ancestry is a more plausible explanation.

**A. Two-way admixture models**

| Target | $Ref_1$ | $Ref_2$ | *P*-value | $Coef_1$ | $Coef_2$ | s.e.m. |
|---|---|---|---|---|---|---|
| Ulsan Korean | Gunsan jar coffin | AR_Xianbei_IA | $2.19\times10^{-1}$ | 0.908 | 0.092 | 0.045 |
| | | Jomon_Ikawazu | $9.51\times10^{-2}$ | 1.004 | -0.004 | 0.007 |
| | | LBK_EN | $1.02\times10^{-1}$ | 1.010 | -0.010 | 0.005 |
| | | Sintashta_MLBA | $1.36\times10^{-1}$ | 1.014 | -0.014 | 0.006 |
| Gunsan jar coffin | Ulsan Korean | AR_Xianbei_IA | $2.23\times10^{-1}$ | 1.101 | -0.101 | 0.054 |
| | | Jomon_Ikawazu | $9.62\times10^{-2}$ | 0.996 | 0.004 | 0.007 |
| | | LBK_EN | $1.03\times10^{-1}$ | 0.990 | 0.010 | 0.005 |
| | | Sintashta_MLBA | $1.38\times10^{-1}$ | 0.986 | 0.014 | 0.006 |

**B. Three-way admixture models**

| Target | $Ref_1$ | $Ref_2$ | $Ref_3$ | *P*-value | $Coef_1$ | $Coef_2$ | $Coef_3$ |
|---|---|---|---|---|---|---|---|
| Ulsan Korean | Gunsan jar coffin | AR_Xianbei_IA | LBK_EN | $5.46\times10^{-1}$ | 0.943<br>(0.048) | 0.072<br>(0.046) | -0.015<br>(0.006) |
| | | AR_Xianbei_IA | Sintashta_MLBA | $6.72\times10^{-1}$ | 0.945<br>(0.048) | 0.074<br>(0.046) | -0.020<br>(0.007) |
| Gunsan jar coffin | Ulsan Korean | AR_Xianbei_IA | LBK_EN | $5.51\times10^{-1}$ | 1.055<br>(0.054) | -0.072<br>(0.053) | 0.017<br>(0.006) |
| | | AR_Xianbei_IA | Sintashta_MLBA | $6.80\times10^{-1}$ | 1.052<br>(0.054) | -0.074<br>(0.052) | 0.022<br>(0.008) |

**Table S5. QpAdm-based admixture modeling of ancient and present-day Koreans and nearby East Asians using distal sources.** (A) We show two-way and three-way admixture model of WLR_BA+Xitoucun and WLR_BA+Xitoucun+Jomon_Ikawazu. We do not detect a significant amount of Jomon contribution in the three-way admixture model. (B) We show three-way admixture model of Miaozigou_MN+Xitoucun+Jomon_Ikawazu. We detect small but significant amount of Jomon contribution in the Gunsan jar coffin individuals and present-day Ulsan Koreans, but it is less suitable than the three-way model using WLR_BA as a northern proxy. WLR_BA, Miaozigou_MN, Xitoucun, and Jomon_Ikawazu represent the estimated ancestry proportion (± 1 s.e.m.) of WLR_BA, Miaozigou_MN, Xitoucun, Jomon_Ikawazu, respectively.

**A. qpAdm modelling results using WLR_BA as a northern proxy**

| Target | *P*-value | WLR_BA | Xitoucun | Jomon_Ikawazu |
|---|---|---|---|---|
| Gunsan jar coffin | $4.59\times10^{-1}$ | 0.898±0.060 | 0.102±0.060 | |
| | $1.14\times10^{-1}$ | 0.877±0.066 | 0.114±0.066 | 0.008±0.013 |
| SGDP Korean | $2.44\times10^{-1}$ | 0.931±0.055 | 0.069±0.055 | |
| | $3.50\times10^{-1}$ | 0.880±0.057 | 0.144±0.058 | -0.024±0.012 |
| Ulsan Korean | $2.20\times10^{-1}$ | 0.952±0.045 | 0.048±0.045 | |
| | $2.07\times10^{-2}$ | 0.921±0.051 | 0.077±0.052 | 0.002±0.010 |
| WLR_LN | $4.89\times10^{-2}$ | 0.852±0.060 | 0.148±0.060 | |
| | $2.97\times10^{-2}$ | 0.758±0.070 | 0.274±0.072 | -0.032±0.013 |
| YR_MN | $1.08\times10^{-1}$ | 0.882±0.055 | 0.118±0.055 | |
| | 6.71×10-2 | 0.822±0.059 | 0.197±0.061 | -0.019±0.012 |
| YR_LN | $1.13\times10^{-1}$ | 0.873±0.048 | 0.127±0.048 | |
| | $1.49\times10^{-2}$ | 0.847±0.055 | 0.162±0.057 | -0.009±0.011 |
| Miaozigou_MN | $1.57\times10^{-1}$ | 0.925±0.090 | 0.075±0.090 | |
| | $1.56\times10^{-1}$ | 0.913±0.104 | 0.118±0.107 | -0.030±0.023 |
| Shimao_LN | $1.75\times10^{-1}$ | 1.056±0.064 | -0.056±0.064 | |
| | $1.11\times10^{-1}$ | 1.021±0.071 | 0.002±0.072 | -0.023±0.015 |
| Upper_YR_LN | $7.69\times10^{-1}$ | 1.021±0.055 | -0.021±0.055 | |
| | $8.91\times10^{-1}$ | 0.983±0.058 | 0.036±0.059 | -0.020±0.012 |

**B. qpAdm modelling results using Miaozigou_MN as a northern proxy**

| Target | *P*-value | Miaozigou_MN | Xitoucun | Jomon_Ikawazu |
|---|---|---|---|---|
| Gunsan jar coffin | $2.53\times10^{-1}$ | 0.908±0.082 | 0.048±0.088 | 0.044±0.018 |
| SGDP Korean | $8.69\times10^{-1}$ | 0.919±0.070 | 0.061±0.073 | 0.020±0.016 |
| Ulsan Korean | $2.49\times10^{-1}$ | 0.940±0.062 | 0.028±0.065 | 0.031±0.014 |
| WLR_BA | $1.56\times10^{-1}$ | 1.096±0.125 | -0.130±0.132 | 0.034±0.025 |
| WLR_LN | $9.30\times10^{-1}$ | 0.873±0.080 | 0.124±0.084 | 0.002±0.018 |
| YR_MN | $2.01\times10^{-1}$ | 0.784±0.076 | 0.214±0.080 | 0.002±0.016 |
| YR_LN | $8.84\times10^{-2}$ | 0.860±0.063 | 0.116±0.066 | 0.024±0.014 |
| Shimao_LN | $5.72\times10^{-1}$ | 1.014±0.082 | -0.018±0.086 | 0.004±0.018 |
| Upper_YR_LN | $3.55\times10^{-1}$ | 1.003±0.075 | -0.016±0.078 | 0.013±0.017 |
